# Distinct antibody-based signatures and functionality distinguish latent and active pediatric tuberculosis

**DOI:** 10.1101/2025.07.21.665984

**Authors:** Nadege Nziza, Wonyeong Jung, Tina Chen, Yixiang Deng, Kees L.M.C. Franken, Tom H.M. Ottenhoff, Sarah Kiguli, Deborah A. Lewinsohn, W. Henry Boom, Harriet Mayanja-Kizza, Mary Nsereko, Sarah M. Fortune, Catherine Stein, Ryan McNamara, Galit Alter, Christina L. Lancioni

## Abstract

**Background:** Tuberculosis (TB), caused by *Mycobacterium tuberculosis* (Mtb), is among the leading causes of death from an infectious agent among children worldwide. Children represent a particularly vulnerable population due to the greater challenges in diagnosis and the higher risk of progression to severe forms of the disease. However, whether different pediatric outcomes relate to distinct immunologic responses remains incompletely understood. Emerging data suggest that Mtb-specific humoral immune responses represent a correlate of protection against Mtb both following natural infection and vaccination.

**Methods:** To determine if immune profiles can distinguish children across the spectrum from Mtb infection to TB disease, as well as children with TB from non-TB lower respiratory tract infection, we mapped the humoral immune response across a panel of 4 dozen Mtb antigens across children presenting with symptoms of active TB (ATB), children with evidence of latent TB infection (LTBI) and children exhibiting non-TB lower respiratory tract infection (non-TB LRTI). Using a custom Luminex assay, Mtb-specific antibody subclass/isotype, Fc receptor (FcR) binding profiles, and functions were profiled across the pediatric groups.

**Findings:** A robust humoral immune response was observed in children with active TB compared to non-TB LRTI, marked by a strong IgA response, that exhibited high FcαR binding. Conversely, children with LTBI uniquely elicited Mtb-specific antibodies with enhanced opsinophagocytic FcγR2A binding, as well as a higher capacity to activate NK cells and neutrophils.

**Interpretation:** There are significant differences in humoral immune profiles across the landscape of pediatric TB, potentially contributing to differential mycobacterial control, and highlighting biomarkers that could guide both diagnostic and therapeutic approaches.

**Funding:** US National Institutes of Health.

## Introduction

Tuberculosis (TB), caused by *Mycobacterium tuberculosis* (Mtb), remains the leading cause of death from an infectious agent globally^1^. Despite widespread Bacille Calmette–Guérin (BCG) vaccination, over one million children under 15 years old develop TB annually, with nearly 250,000 resulting deaths. Children under five years old are particularly vulnerable, facing higher risks of disease progression, morbidity, and mortality^2,3^. While TB progresses in approximately 5-10% of adults, Mtb-infection leads to pulmonary disease within two years of exposure in nearly 20% of children under 5 years old and is more likely to lead to severe, disseminated forms of TB during early childhood^3^. Understanding TB pathophysiology in children is critical for improving diagnostics, identifying therapeutic targets, and advancing next-generation vaccines.

Across several infectious diseases, there is mounting data to support that differences in antibody responses can reflect distinct disease states^4-8^. Recent studies have shown that the quality of Mtb-specific antibody responses—including isotype, subclass, and Fc-glycosylation—varies between adults with active TB (ATB) and asymptomatic and/or latent TB infection (LTBI)^7,9-12^, and can predict progression^13^ and TB recurrence^14^. These findings among adults suggest that Mtb-specific antibody responses could have a role as a diagnostic tool to identify young children with LTBI and ATB. There is also emerging evidence that Mtb-specific antibodies may offer protection against progression to ATB following infection. For example, Mtb-specific antibodies are linked to intracellular control of infection^11^, and passively transferred antibodies reduce TB disease severity and offer protection against lethal challenge in animal models^15,16^. Mtb-specific antibodies are also associated with near-sterilizing protection against Mtb-infection following intravenous BCG vaccination^6^ in non-human primates. However, the immune mechanisms regulating the delicate balance between host control and disease in children are not understood, and identifying correlates of protective immunity remains challenging. Understanding the specificity of a protective humoral immune response may be key to development of a next-generation vaccine that effectively protects all young children from TB. In addition, the development of a Mtb-specific humoral profile that accurately distinguishes young children with lower-respiratory tract infections (LRTI) due to TB from other common bacterial etiologies, would provide a much needed diagnostic tool to facilitate initiation of appropriate clinical treatment.

Here we utilized a broad array of previously identified immunogenic Mtb-antigens (including surface, secreted, and intra-cellular antigens) to address two fundamental questions: first, can differences in Mtb-specific functional antibody profiles distinguish between ill young children with TB and non-TB LRTI, as well between children with ATB from LTBI; and second, do differences in Mtb-specific functional antibody profiles offer insight into the immunopathology of pediatric Mtb-infection and TB disease.

## Material and methods

### Study subjects

Children 1–60 months old admitted with severe pneumonia were recruited from Mulago Hospital in Kampala, Uganda, from 2011-2014^17^. All children underwent standardized diagnostic evaluation for pulmonary TB; additional studies for disseminated TB were performed if clinically indicated. Children with a history of TB, and children on TB treatment for seven or more days before recruitment, were excluded. All children were tested for HIV using a rapid-diagnostic assay; infants under 18 months were tested for HIV via PCR. Children-living-with-HIV were not receiving antiretroviral therapy (ART). All children underwent a single blood draw prior to initiation of TB treatment for isolation and cryopreservation of plasma.

Children were evaluated 2 months following hospital discharge to assess response to therapy and subsequently classified as having: microbiologically confirmed TB (pulmonary and/or disseminated TB; ATB) or non-TB lower-respiratory-tract-infection (non-TB LRTI), by consensus review by three pediatricians and using standardized definitions. ATB was defined by isolation of Mtb from an induced sputum sample by acid-fast bacillus (AFB) culture in a child with signs/symptoms of pulmonary and/or disseminated TB disease, as defined by: chest X-ray (CXR) consistent with pulmonary TB; clinical signs/symptoms suggestive of TB including persistent cough of 2 weeks duration or longer, failure to thrive, unexplained fever for one week or longer, persistent lethargy, known close contact with adult with confirmed TB, positive tuberculin skin test (TST), or clinical response to TB treatment. Non-TB LRTI was defined by all of the following: hospitalized with signs and symptoms of pneumonia; absence of bacteriologic confirmation of TB, child did not demonstrate two or more features of TB disease, negative TST, no known TB exposure, and clinical improvement in absence of TB treatment.

Healthy Ugandan children, 1–60 months old, who were a household contact of an adult with confirmed TB, were recruited in Kampala (2002-2012)^18^. Cryopreserved plasma obtained at enrollment from children with a negative diagnostic evaluation for TB disease but positive TST, and who remained asymptomatic following two years of observation, were included. These children were classified as LTBI.

The study was approved in Uganda and U.S. Written informed consent was obtained from parents/guardians in their preferred language. Key demographic characteristics are detailed in **Supplemental Table 1**.

### Antigens

Forty-seven Mtb antigens and six nonTB antigens were studied. Mtb antigens included purified lipoarabinomannan (LAM) (BEI), and purified protein derivative (PPD) (Statens Serum Institute). The remaining 45 Mtb proteins were recombinantly produced and provided by Dr. Tom Ottenhoff and Kees Franken (**Supplemental Table 2**)^19^. Antigen selection was based on immunogenic properties^7,19-21^.

### Antibody isotype, FcR binding, and functions

Antibody isotype and subclasses (IgG1, IgG2, IgG3, IgG4, IgM, and IgA1), as well as Fc-receptor (FcR) binding profiles (FcyR2A, FcyR2B, FcyR3A, FcyR3B, and FcαR) were measured using a custom Luminex assay^22^. Plasma samples were diluted between 1:50 and 1:500, depending on the secondary reagent. For functional analyses, antibody-dependent complement deposition (ADCD), cellular phagocytosis (ADCP), neutrophil phagocytosis (ADNP), and antibody-dependent NK activation (ADNKA) were performed^23,24,25^ with 1:20 diluted plasma samples. MFI values were analyzed on an iQue analyzer (Intellicyt).

### Statistical analysis

GraphPad Prism (v·9·2·0) and RStudio (v·1·3 and R v·4·0) were used for analyses. Multivariate analyses compared nonTB-LRTI and TB groups (ATB and LTBI), as well as ATB and LTBI. Data were normalized followed by a least absolute shrinkage and selection operator (LASSO) approach for feature selection. Partial least square discriminant analysis (PLS-DA) models were performed using LASSO-selected features, for classification and visualization, followed by a ten-fold cross-validation to assess model accuracy.

### Linear model adjusting for age

To evaluate the significance of the association between TB status and systems serology features while controlling for age, we fitted a linear model that included TB status and confounders including age, and a nested linear model that only included the confounder:

‐ Null model: systems serology measurement ∼ age
‐ Full model: systems serology measurement ∼ TB status + age

A likelihood ratio test was performed to assess whether the full model provided a significantly better fit to the systems serology data compared to the null model. The group coefficient from the full model and the p-value from the likelihood ratio test—adjusted for multiple comparisons using the Benjamini-Hochberg method—were visualized in a volcano plot.

## Results

### TB disease state and Mtb-specific humoral immunity

Here we broadly profiled humoral immune responses among cohorts of children under 60 months living in urban Uganda, a high-burden TB setting. Cohorts included children with active TB (ATB; hospitalized with confirmed pulmonary or disseminated TB; n=56), children hospitalized with non-TB LRTI (n=62), and TB-exposed children with latent TB infection (LTBI; n=14). Mtb-specific humoral immune responses were profiled across a panel of 47 Mtb-derived antigens previously defined as immunogenic in adults^7,26^. A custom Luminex assay was used to analyze Mtb-specific IgG1, IgG2, IgG3, IgM, IgA1, and Fc-receptor binding profiles (FcγR2A, FcγR2B, FcγR3A, FcγR3B, FcαR). The Mtb-specific humoral profile was plotted across all children with ATB, LTBI, and non-TB LRTI, in a blinded fashion (**Figure 1A**). Mtb-specific IgG1 and IgM responses were heterogenous, with responses detected in all three groups across different antigens. Slight enrichment of Mtb-specific IgG1 and IgM responses was detected in children with ATB. Conversely, Mtb-specific IgG2 and IgG3 responses were highest in a subgroup of children with ATB, weakly observed in children with LTBI (**Figure 1A, B**), and were largely negative in children with non-TB LRTI. Mtb-specific IgA exhibited the most distinct profile, with high levels across nearly all antigens in children with ATB, moderate levels in children with LTBI, and sporadic responses in children with non-TB LRTI. Antibody levels across groups were compared for a representative Mtb antigen, Ag85A/B, and a non-Mtb antigen (Influenza HA/Flu) (**Figure 1B**). Ag85A/B-specific IgG1 and IgA responses were highest in children with ATB and lower in children with LTBI, but both were higher than levels observed in non-TB LRTI (**Figure 1A, B**). Conversely, Flu-responses were comparable across all groups. IgG1 and IgA responses were higher in older children, with responses emerging in the first 6-12 months of life in children with ATB. Similarly to IgA, FcαR –a receptor specific of IgA– showed strong binding levels to Mtb-specific antibodies within the ATB population, particularly in older children (**Supplemental Figure 2**). Mtb-specific antibody binding to FcγR2A was particularly elevated in LTBI, while FcγR2B, FcγR3A and FcγR3B binding exhibited responses similar to IgGs, with an enrichment in TB groups.

**FIGURE 1.**
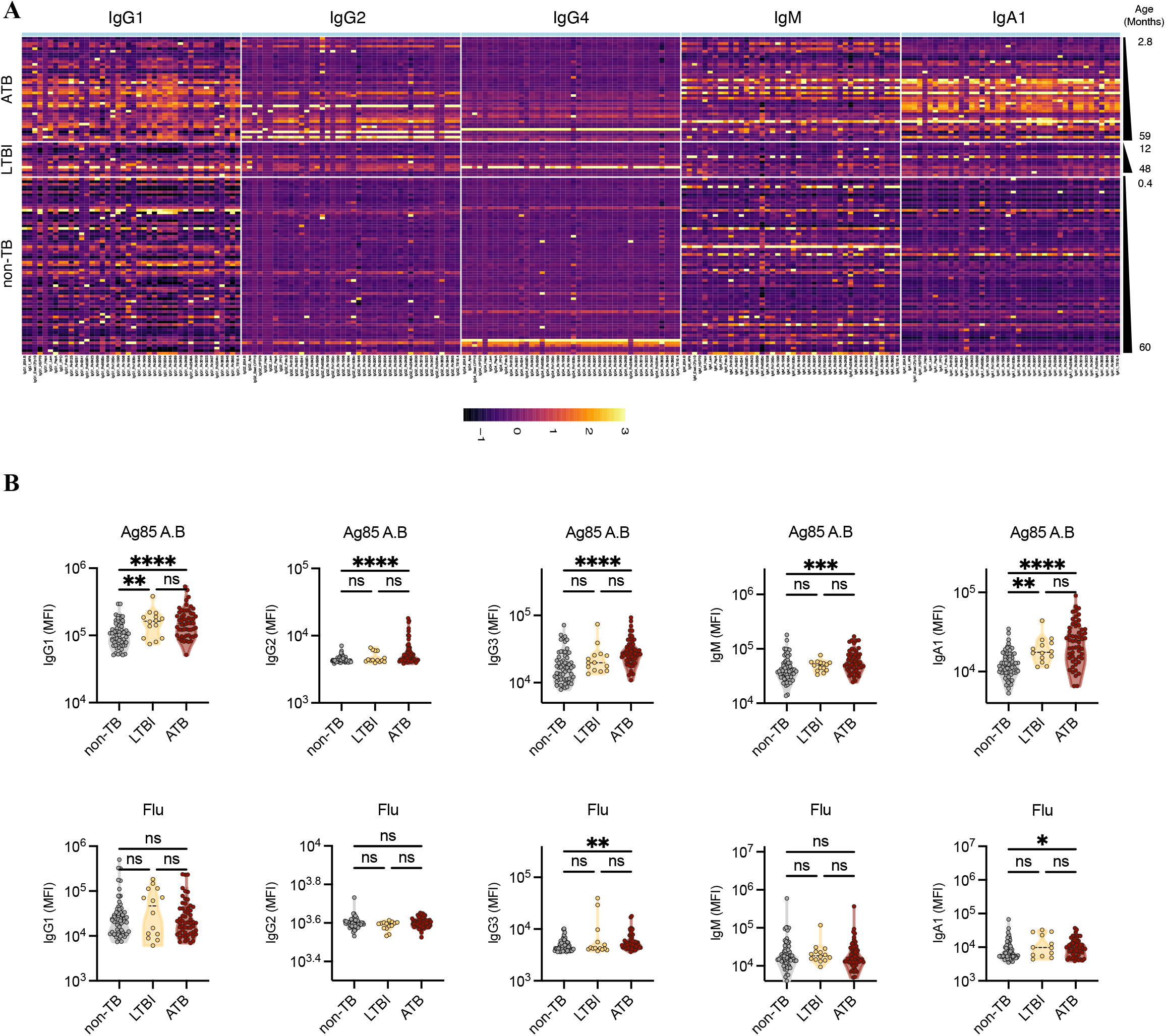
Mtb-specific humoral response across pediatric cohorts. Relative levels of antibodies were measured via Luminex in the plasma of ATB, LBTI and nonTB individuals aged 0·4 to 60 months. (A) Heatmap illustrate IgG1, IgG2, IgG4, IgM, and IgA1 levels for each individual ordered by age. MFI data were Z-scored across columns. Each column represents one antigen-specific antibody feature, each row indicates a different individual. (B) The dot plots show the univariate analysis of antibody levels against Ag85 A and B, as well as Flu. Significance was calculated by using two-sided Mann-Whitney U-test, *p < 0·05, **p < 0·01, ***p < 0·001, ****p < 0·0001.

### Antibody levels and FcR binding reveals strong Mtb responses in pediatric TB

We next defined features of the Mtb-specific humoral immune response that could discriminate between TB and non-TB disease states. We expanded our profiling of Mtb-specific humoral immune responses to other isotypes and Fc-receptor binding antibodies (**Figure 2**). Using this broader dataset, we first defined whether a specific set of Mtb-specific antigens could discriminate TB cases (including both ATB and LTBI) from non-TB LRTI cases in children aged 12-60 months, after waning of maternally transferred antibodies (**Figure 2A**). Mtb-specific humoral immune responses were selectively elevated in children from the TB compared to non-TB LRTI groups (**Figure 2**), as well as the ATB relative to non-TB LRTI groups (**Supplemental Figure 2**). More specifically, Mtb-specific IgG1 and IgG2 levels, along with strong FcγR2A, FcγR2B, and FcγR3A binding, reflected signatures of Mtb-infection targeting a range of antigens. Similar results were seen for IgG3 and FcγR3B (**Supplemental Figure 3**). Notably, Mtb-specific IgA1/FcαR levels were the most significantly differentially expanded subclass in TB compared to non-TB cases (**Figure 2A**). These findings highlight the potential of Mtb-specific antibody binding profiles to serve as a diagnostic tool to distinguish young children with pulmonary and disseminated TB from children with non-TB LRTI.

**FIGURE 2.**
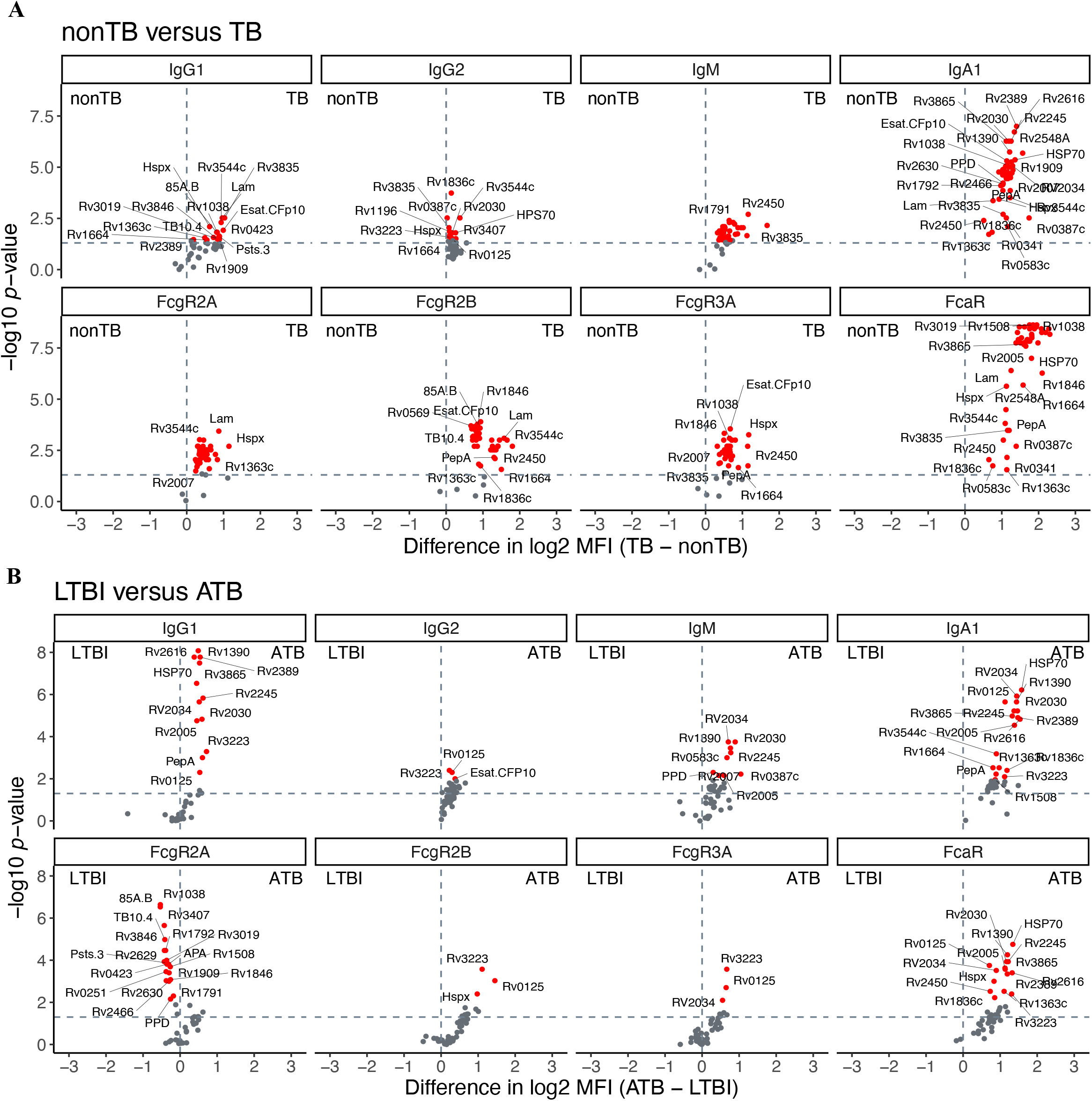
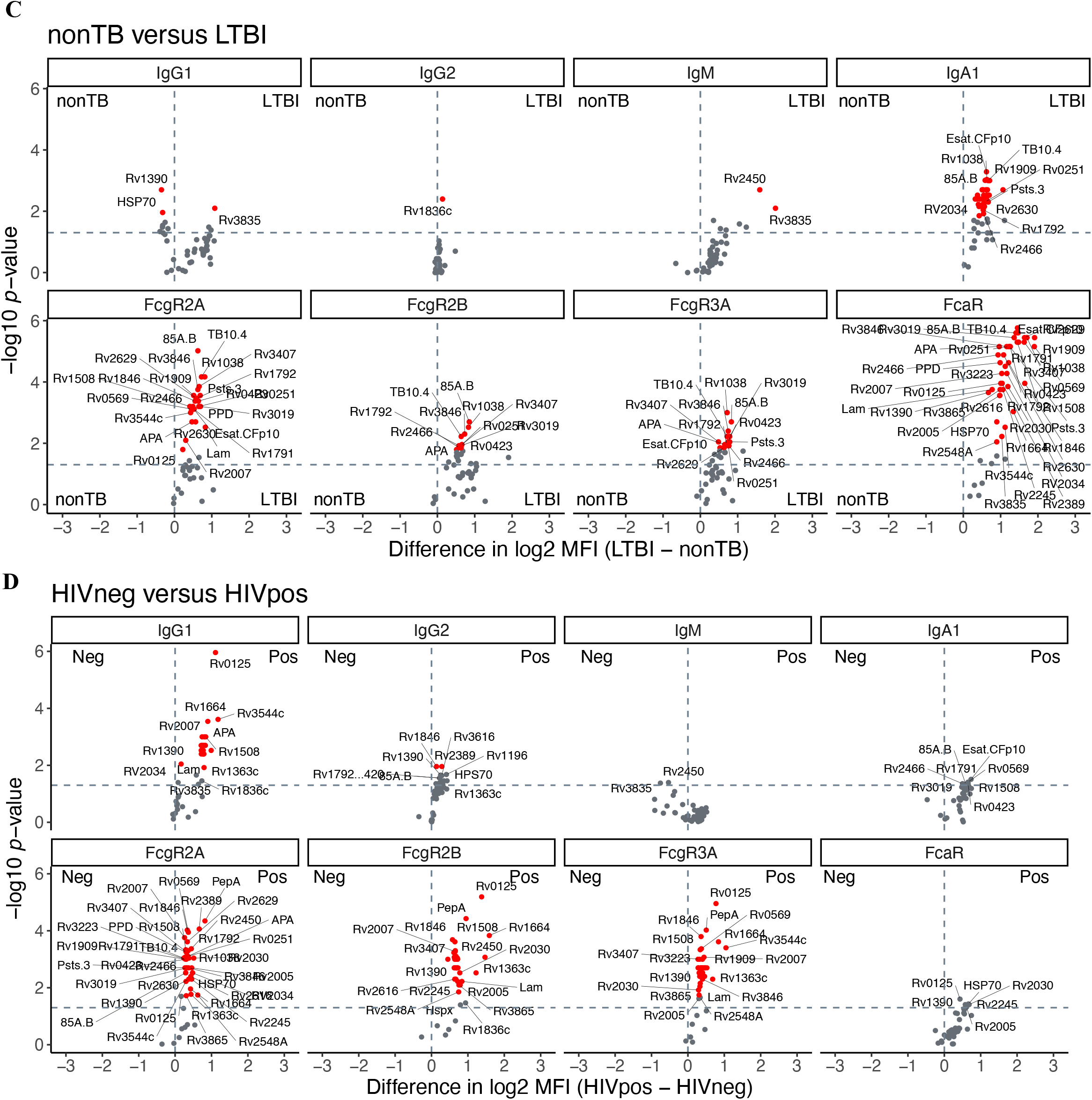
Tuberculosis is associated with strong Mtb-specific antibody levels and FcR binding. Relative levels of IgG1, IgG2, IgM and IgA1, as well as FcγR2A, FcγR2B, FcγR3A and FcαR binding against 47 Mtb antigens were quantified via Luminex in the plasma of children aged 12-60 months, comparing nonTB and TB (including ATB and LTBI) individuals (A), ATB and LTBI groups (B), nonTB and LTBI groups (C), as well as ATB HIV- and ATB HIV+ (D). The volcano plots characterize the magnitude (log2 fold change of TB/nonTB, ATB/LTBI, and nonTB/LTBI) and the significance (p values) of antibody levels between groups. Values above black dashed lines are statistically different between the groups (p < 0·05). For adjusted p-values, significant data are shown in red, non-significant differences are in grey.

We next determined whether particular Mtb-antibody profiles could discriminate children with ATB from those with LTBI (**Figure 2B**). Children with ATB exhibited significant expansion of Mtb-antibody responses, with IgA1/FcαR responses most expanded in ATB compared to LTBI. Conversely, Mtb-specific FcγR2A binding levels were the sole set of antibody features selectively expanded in children with LTBI compared to ATB. Importantly, no significant differences were noted in Mtb-specific antibody profiles across BCG vaccinated or unvaccinated children (**Supplemental Figure 4A**), across TST-positivity (**Supplemental Figure 4B**), or sex (**Supplemental Figure 4C**). These data highlight the potential use of Mtb-specific antibody binding profiles as biomarkers of ATB and LTBI in young children.

Comparison of children with LTBI and nonTB-LRTI revealed strong differences in IgA1, FcγR2A, FcγR2B, FcγR3A, and FcαR binding levels, with stronger responses in children with LTBI (**Figure 2C**). Slightly higher IgM levels were identified in children with LTBI compared to nonTB-LRTI, while IgG1 was enriched in both groups, depending on the Mtb antigen.

HIV modulates antibody profiles in Mtb-infected adults^7,26,27^. We examined antibody profiles across children with ATB who were living-with and without-HIV. (**Figure 2D**). Living-with-HIV was associated with higher Mtb-specific IgG, FcγR2A, FcγR2B, and FcγR3A binding levels. In contrast, no differences were observed in Mtb-specific IgG2, IgM, or FcαR binding levels, suggesting that some features are not affected by HIV status.

### Multivariate analysis identifies biomarkers that distinguish TB and non-TB pediatric cases

We next determined if a minimal set of Mtb-specific antibody features could discriminate between children aged 12-60 months with TB (ATB and LTBI) and non-TB LRTI. All data were integrated and a LASSO feature down-selection followed by PLS-DA classification applied (**Figure 3A and B**). Mtb-specific antibody profiles differed across the 2 groups, differentiated along latent variable 1 (LV1) (**Figure 3A**). Children in the TB group displayed a distinct Mtb-specific antibody profile in multivariate space. Of the total 506 antigen-specific antibody features included in analysis, only six features were sufficient to separate TB and non-TB antibody profiles (**Figure 3A and B**), with Mtb-specific antibody features all enriched among children in the TB group. An area under the receiver operating characteristic (ROC) curve (AUC) demonstrated 0·945 (0·890-1·000) classification accuracy using the six LASSO-selected antibody features (**Figure 3C)**. Across the LASSO signature, Rv2034 and Esat6/CFP10 FcαR responses, and Rv1846 IgA responses were all significantly and preferentially enriched in children in the TB group (**Figure 3D)**. Additionally, these children exhibited elevated Rv1038-specific FcγR2B binding antibodies, Esat6/CFP10 FcγR3A binding antibodies, and elevated Rv2007-specific IgG3 levels. Finally, a feature clustering regression was performed to define whether particular feature classes were preferentially enriched among children in the TB group (**Figure 3E**). Across isotypes and subclasses, in addition to ESAT6/CFP10 IgG1 and IgG3 titers, nearly all Mtb-specific IgA responses were enriched among children in the TB group. Likewise, across Fc-receptor binding features, FcαR binding antibodies were among the most highly enriched features in children within the TB group, followed by a subset of Mtb-specific FcγR3b-binding antibodies.

**FIGURE 3.**
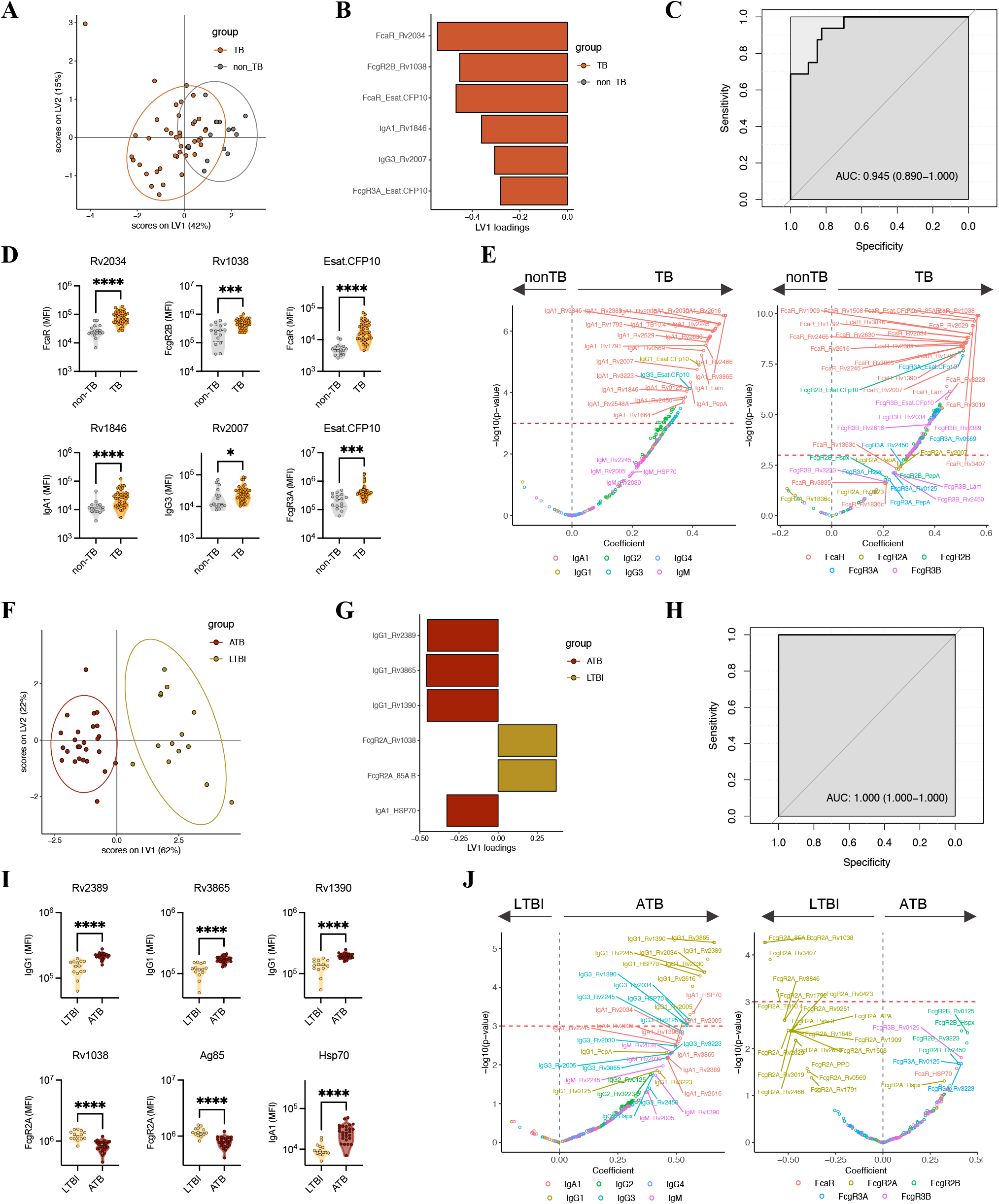
Ant Antibody-based signatures discriminate between children with and without TB, and between LTBI and active TB. Antibody levels and FcR binding were measured in the plasma of children aged 12-60 months with TB versus nonTB (A-E) in addition to ATB versus LTBI (F-J). Multivariate analysis comparing antibody response in TB and nonTB groups (A, B), as well as ATB and LTBI (F, G). (A, F) Score plot from PLS-DA model with each dot representing each individual and x- and y-coordinates showing latent variable 1 (LV1) and latent variable 2 (LV2), respectively. Ellipses show 95% confidence intervals. PLDS-DA models were trained using LASSO-selected features that are plotted on VIP plots (B, G). The colors of the bars are indicating the group in which the features have the highest mean. (C, H) ROC curves showing the sensitivity and the specificity of potential biomarkers discriminating TB and nonTB groups (C), as well as ATB and LTBI (H). (D, I) Dot plots representing the univariate analysis of the LASSO-selected features. Significance was calculated by using two-sided Mann-Whitney U-test, *p < 0·05, ***p < 0·001, ****p < 0·0001. (E, J) Volcano plots illustrating the results from linear regression for the difference between TB and nonTB (E), as well as ATB and LTBI (J), adjusting for age. Features above horizontal dashed lines are significantly enriched in one group, with p-value <0·05 after multiple test corrections by Benjamini-Hochberg procedure.

### Multivariate analysis identifies biomarkers that distinguish pediatric LTBI and ATB

While the TST and interferon-gamma-release-assays (IGRA) cannot distinguish between ATB and LTBI, emerging data point to significant differences in Mtb-specific antibody profiles across adult LTBI and ATB^7,11,27,28^. Whether similar signatures can be defined in pediatric TB disease versus LTBI remains uncertain. Thus, we next compared Mtb-specific antibody features across children with ATB and LTBI (**Figure 3F-J**). The PLS-DA classification using LASSO-selected features demonstrated that a combination of six Mtb-specific antibody features (out of 506 analyzed), was sufficient to separate children with ATB from those with LTBI (**Figure 3F and G**). IgG1 levels to Rv2389, Rv3856, Rv1390, and HSP70-specific IgA1 levels were all selectively expanded in children with ATB compared to LTBI. Conversely, Rv1038- and Ag85 A/B-specific FcγR2A binding antibody levels were higher in children with LTBI (**Figure 3G**). The combination of these markers provided perfect prediction, an AUC of 1 (1·000-1·000) (**Figure 3H**). All LASSO-selected features exhibited univariate significance (**Figure 3I**). Finally, feature clustered regression analysis with adjustment for age highlighted enrichment of Mtb-specific titers among children with ATB, but a selective expansion of particular Mtb-specific FcγR2A-binding antibody responses in children with LTBI.

These data underscore the potential for Mtb-specific antibody biomarkers to discriminate between pediatric ATB and LTBI, as well as ATB and non-TB LRTI, among young children living in a TB-endemic setting.

### Latency is associated with strong functional antibody-mediated activation of immune cells

To evaluate whether differences in overall antibody levels and binding profiles were associated with differences in antibody effector functions across TB disease states, we analyzed antibody-dependent cellular activation, including ADNP (**Figure 4A and B**), ADCP (**Figure 4C and D**), ADNKA (with %CD107a) (**Figure 4E and F**), and ADCD (**Figure 4G and H**) for Lam, Hsp70, Rv2034, and Rv2007. ADNP, ADCP, and ADNKA activity was similar between children in TB (ATB and LTBI) and non-TB LRTI groups (**Figure 4 A, C, and E**). One exception was Rv2034, where antibodies from children with non-TB LRTI demonstrated increased ADCP activity. Conversely, high Lam-, HSP70-, and Rv2034-specific ADNP, as well as Lam-, Rv2007-specific ADCP, and Rv2007-specific ADNKA were detected in children with LTBI compared to children with ATB (**Figure 4B, D, and F)**. These findings may point to a role for particular antigen-specific antibody functions in controlled infection. Regarding ADCD, a higher response was observed in the TB group compared to the non-TB LRTI group for Rv2007 (**Figure 4G**), whereas similar levels were observed between ATB and LTBI (**Figure 4H**). Finally, antibody-mediated functionality for non-TB antigens, including influenza and tetanus, showed similar responses between the TB and non-TB LRTI groups (**Supplemental Figure 5**). Correlation heatmaps, examining the relationship of antibody features between cohorts, revealed negative correlations across LTBI when compared to either ATB or non-TB LTBI, particularly for opsinophagocytic antibody functions (**Figure 4I**). These results underscore the selective enrichment of antibody effector functions among LTBI and highlight two key differences in Mtb-specific humoral immunity in early life: 1) preferential elevation of select functional antibodies in LTBI; and 2) preferential elevation of Mtb-specific binding antibodies in ATB.

**FIGURE 4.**
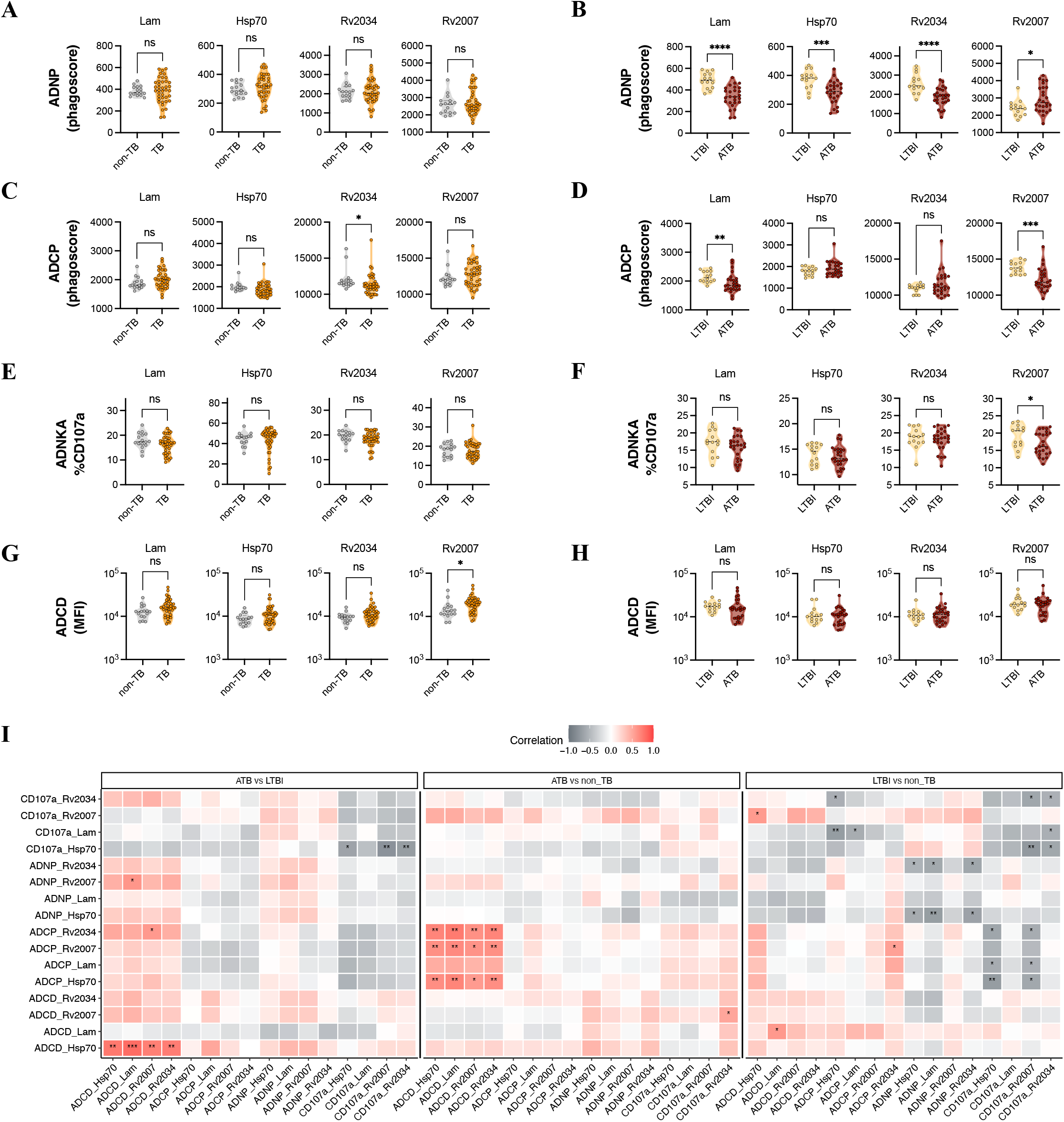
Analysis of antibody functionality among children with and without TB. Antibody-dependent complement deposition (ADCD), Antibody-dependent cellular phagocytosis (ADCP), Antibody-dependent neutrophil phagocytosis (ADNP), and Antibody-dependent NK cell activation (ADNKA) with %CD107a against TB antigens were analyzed in the plasma of children aged 12-60 months. (A-H) The dot plots show differences in antibody functions between TB and nonTB (B, D, F, H), as well as ATB and LTBI (C, E, G, I). Significance was calculated by using two-sided Mann-Whitney U-test, *p < 0·05, ***p < 0·001, ****p < 0·0001. (I) Heatmaps show the correlation between antibody response in ATB versus LTBI; ATB versus non-TB; LTBI versus nonTB. Spearman correlations were used to calculate the level of correlation between the different groups, with positive correlations in red and negative correlations in grey. Statistical differences were calculated using non-parametric two-sided Mann-Whitney U test and corrected by the BH method. *p < 0·05, **p < 0·01, ***p < 0·001.

## Discussion

Compared to adults, children are more susceptible to infections by various pathogens, including Mtb, which leads to significant morbidity and mortality^3^. Although precise mechanisms driving vulnerability to TB during early life are not well defined, the immature state of both the innate and adaptive immune responses likely contributes to early disease progression following Mtb-exposure^2^. Moreover, unique aspects of the pathophysiology and clinical presentation of pediatric TB make establishing the diagnosis challenging and often lead to treatment delays. Specifically, the lack of non-sputum based rapid and specific point-of-care diagnostic tools has hindered TB control in children^29^ and distinguishing young children with LRTI secondary to TB from other bacterial causes in TB-endemic settings remains challenging. Plasma antibody-based tests offer promising alternatives to current diagnostic methods that cannot distinguish LTBI from TB disease^7^. However, little is known about Mtb-specific humoral immunity in children; previous studies using strictly quantitative measurements of canonical antigen-specific antibody levels have been unsuccessful in identifying Mtb-specific humoral biomarkers for TB diagnosis in children. Therefore, we explored both qualitative differences in humoral immune profiles and the breadth of antigen-specific antibody responses across acutely ill Ugandan children with TB and non-TB LRTI, and explored the potential of antibodies to distinguish pediatric ATB from LTBI.

A custom Luminex assay was used to analyze antibody subtypes and functionality against 47 Mtb antigens. We observed a robust IgA/FcαR response associated with pediatric Mtb-infection and TB disease. This IgA-signature was unexpected, as IgA responses are generally reported to develop later in childhood, unlike IgG1 and IgG3 responses that develop first during infancy, followed by IgG2 and IgG4^30^. Higher IgA levels in children with ATB may be related to elevated antigen burden within the respiratory mucosa, resulting in enhanced IgA class-switching. These data point to the high discriminatory potential of the IgA isotype, which we have found to be specifically produced in Mtb-infected children and less likely to be detected in young children with non-TB LRTI despite living in a TB-endemic region with high rates of infant BCG-vaccination. Furthermore, particular Mtb-specific IgG2, IgG3, and IgM levels, as well as FcγR2A, FcγR2B, FcγR3A, and FcγR3B binding levels, contributed to discrimination between children with TB and non-TB LRTI. These results highlight the high specificity and sensitivity of antibody-based biomarkers and their potential for TB diagnostic approaches for young children hospitalized with pneumonia in TB-endemic settings.

Identification of children with latent, or asymptomatic Mtb is critical for initiating TB preventive therapy and could contribute to reducing TB-related morbidities and mortality. Although TST and IGRA testing can assess for immune sensitization to Mtb as a proxy for infection, both assays have limited performance in young children and neither can distinguish between LTBI and ATB. Our data comparing ATB and LTBI highlighted significant differentiation between the two groups using Mtb-specific antibody profiling, potentially providing a path to improve LTBI diagnosis. Specifically, Mtb-specific IgG1 levels were enriched in both ATB and LTBI cases, while IgA and FcαR remained the strongest signatures of ATB disease, followed by FcγR2A, FcγR2B, and FcγR3A binding. These results could guide the development of diagnostic tools for LTBI, which are critically needed to improve clinical care and prevent TB progression.

The maturation of the humoral immune system in infants and young children leads to age-dependent differences in antibody responses to various antigens. Given that children under 5 years of age are at the highest risk of progressing to ATB and disseminated TB^2^, it is crucial to define biomarkers tailored in an age appropriate manner. Here we demonstrate that after adjusting for age in a linear regression model, strong age-independent differences in Mtb-specific antibody profiles persisted between cohorts of children.

Our study does include limitations. First, only Ugandan children were included and our findings need to be validated in diverse populations to establish geographical generalizability. Only a limited number of functional assays were performed and we were unable to assess if Mtb-specific antibodies from children with LTBI enhanced antimicrobial control. None of our children-living-with-HIV were receiving ART at enrollment, thus potentially limiting relevance to populations of children on ART.

An effective immune-based diagnostic approach to pediatric TB is urgently needed to optimize clinical care, establish correlates of immunity for vaccine trials, and strengthen studies of TB epidemiology in young children. Here we identified a multiparametric Mtb-specific antibody-based signature that distinguishes between acutely ill children with and without TB LRTI, as well as between children with LTBI and ATB. Our identification of an expanded Mtb-specific functional antibody profile in children with LTBI suggests that the role of humoral immunity in protection against pediatric TB should be prioritized for further study.

## Contributors

All authors have final responsibility for the decision to submit for publication. C.L.L., G.A., N.N., R.M., C.S. and S.M.F. contributed to the conception and/or interpretation of the work. N.N., G.A., R.M., W.J., T.C., Y.D., K.F, T.O., contributed to the laboratory protocols, statistical analysis, and developed the figures and tables that were later reviewed by all the authors. S.K., D.A.L., W.H.B., M.N., and C.L.L. contributed to development of human subjects protocols, acquisition and interpretation of clinical data, and establishment of clinical cohorts. N.N., C.L.L. and G.A. wrote the first manuscript that was later reviewed by all the authors. The final version of the manuscript was reviewed by all authors. N.N., G.A., and R.M. accessed and verified all the data. All authors had access to all the data.

## Data Sharing

De-identified data used in this study are available upon request to the corresponding author.

## Acknowledgements

This work was supported by the NIH 1R01 AI157807-01A1 (CLL/CS); NIH Contract no. HHSN272200900053C (DAL); CWRU Tuberculosis Research Unit contract HHSN266200700022C (WHB). We acknowledge support from the IMPAcTB Consortium, Terry and Susan Ragon, and Mark and Lisa Schwartz.

