## Supplemental materials for "Distinct antibody-based signatures and functionality distinguish latent and active pediatric tuberculosis"

**<u>Table of Contents:</u>**


| | |
|---|---|
| **Supplemental Table 1** | **p. 2** |
| **Supplemental Table 2** | **p. 3, 4** |
| **Supplemental Figure Legends** | **p. 5** |
| **Supplemental Figure 1** | **p. 6** |
| **Supplemental Figure 2** | **p. 7** |
| **Supplemental Figure 3** | **p. 8** |
| **Supplemental Figure 4** | **p. 9,10** |
| **Supplemental Figure 5** | **p. 11** |

**Supplemental Table 1. Study participant demographic and clinical findings**

| | Confirmed Active TB (n=56) Pulmonary (n=48) | Disseminated (n=8) | LTBI (n=14*) | Non-TB LRTI (n=62) |
|---|---|---|---|---|
| Age (months; mean±SD) | 21·4 ±15·5 | 23·5±16·5 | 32·3±15·9 | 20·1±12·3 |
| Sex (% female) | 54% | 50% | 38.4% | 37% |
| TST positive (%) | 62·5% | 12·5% | 100% | 0% |
| BCG Vaccinated (%) | 70·8% | 62·5% | 100% | 80·6% |
| Living-with-HIV | 29·1% | 25% | 0% | 0% |

*Complete demographic information not available for 1 child with LTBI

**Supplemental Table 2. Mycobacterial antigens tested**

| H37Rv code | Alternative name | Function |
|---|---|---|
| Rv0125 | pepA | Probable serine protease PepA (serine proteinase) (MTB32A) |
| Rv0251 | hsp20, hrpA, acr2 | heat shock protein |
| Rv0288 | esxH | protein TB10.4 |
| Rv0341 | iniB | Isoniazid inducible gene protein IniB |
| Rv0350 | dnaK, hsp70 | heat shock protein |
| Rv0387c | | Conserved hypothetical protein |
| Rv0423 | thiC | thiamine biosynthesis |
| Rv0569 | | conserved protein |
| Rv0583c | lpqN | Probable conserved lipoprotein LpqN |
| Rv0928 | Psts.3 | phosphate uptake |
| Rv1038 | esxJ | ESAT-6 like protein EsxJ |
| Rv1196 | PPE18 | PPE family protein PPE18 |
| Rv1363c | | Possible membrane protein |
| Rv1390 | rpoZ | predicted RNA polymerase |
| Rv1508 | Rv1508c | Probable membrane protein |
| Rv1664 | pks9 | Probable polyketide synthase Pks9 |
| Rv1791 | PE19 | PE family protein |
| Rv1792 | esxM | ESAT-6 like protein |
| Rv1836c | | Conserved protein |
| Rv1846 | blaI | transcriptional repressor |
| Rv1860 | APA | Alanine and proline rich secreted protein Apa (fibronectin attachment protein) (immunogenic protein MPT32) (antigen MPT-32) (45-kDa glycoprotein) (45/47 kDa antigen) |
| Rv1886 | fbpB, mpt59, 85B | lipid metabolism |
| Rv1909 | furA | Iron metabolism |
| Rv2005 | | stress response family protein |
| Rv2007 | fdxA | Iron metabolism |
| Rv2030 | | conserved protein |
| Rv2031c | Hspx | Heat shock protein HspX (alpha-crystallin homolog) (14 kDa antigen) (HSP16.3) |
| Rv2034 | | ArsR repressor protein |
| Rv2245 | kasA | lipid metabolism |
| Rv2389 | rpfD | predicted resuscitation-promoting factor |
| Rv2450 | rpfE | predicted resuscitation-promoting factor |
| Rv2466 | | conserved protein |
| Rv2548A | | Conserved protein |
| Rv2629 | | conserved protein |
| Rv2630 | | conserved protein |
| Rv3019 | esxR, TB10.3 | ESAT-6 like protein |
| Rv3223 | sigH, rpoE | RNA polymerase sigma factor |
| Rv3407 | vapB47 | predicted antitoxin |
| Rv3544c | fadE28 | Probable acyl-CoA dehydrogenase FadE28 |
| Rv3616 | espA | ESX-1 secretion associated protein |
| Rv3775.Rv3874 | ESAT6/CFP10 | fusion protein |
| Rv3804 | fbpA, mpt44, 85A | lipid metabolism |
| Rv3835 | | Conserved membrane protein |
| Rv3846 | sodA, sodB, sod | superoxide dismutase |

| | | |
|---|---|---|
| Rv3865 | espF | ESX-1 secretion associated protein |

**Supplemental Figure 1. Mtb-specific FcR binding in ATB, LTBI and non-TB groups.** FcR binding was measured via Luminex in the plasma of ATB, LBTI and nonTB individuals aged 0.4 to 60 months. (1) Heatmap illustrate FcγR2A, FcγR2B, FcγR3A, FcγR3B and FcαR binding levels for each individual ordered by age. MFI data were Z-scored across columns. Each column represents one antigen-specific antibody feature, each row indicates a different individual. (B) The dot plots show the univariate analysis of FcR binding levels against Ag85 A and B, as well as Flu. Significance was calculated by using two-sided Mann-Whitney U-test, *$p < 0{\cdot}05$, **$p < 0{\cdot}01$, ***$p < 0{\cdot}001$, ****$p < 0{\cdot}0001$.

**Supplemental Figure 2. Strong Mtb-specific antibody levels and FcR binding in ATB.** Relative levels of IgG1, IgG2, IgM and IgA1, as well as FcγR2A, FcγR2B, FcγR3A and FcαR binding against 47 Mtb antigens were quantified via Luminex in nonTB and ATB children aged 12-60 months. The volcano plots characterize the magnitude (log2 fold change of ATB/nonTB) and the significance (p-values) of antibody levels between groups. Values above black dashed lines are statistically different between the groups ($p < 0{\cdot}05$). For adjusted p-values, significant data are shown in red, non-significant differences are in grey.

**Supplemental Figure 3. IgG3 and FcgR3B responses against Mtb antigens.** Relative levels of IgG3 as well as FcγR3B binding against 47 Mtb antigens were quantified via Luminex in the plasma of children aged 12-60 months. (A) Comparison between nonTB and TB individuals. (B) Comparison between LTBI and ATB individuals. (C) Comparison between nonTB and LTBI individuals. (D) Comparison between nonTB and ATB individuals. The volcano plots characterize the magnitude (log2 fold change) and the significance (p values) of antibody response between groups. Values above black dashed lines are statistically different between the groups ($p < 0{\cdot}05$). For adjusted p values, significant data are shown in red, non-significant differences are in grey.

**Supplemental Figure 4. Impact of HIV infection, BCG vaccination, TST, and sex on antibody response in ATB.** Relative levels of IgG1, IgG2, IgM and IgA1, as well as FcγR2A, FcγR2B, FcγR3A and FcaR binding against 47 Mtb antigens were quantified via Luminex in the plasma of ATB individuals aged 12-60 months. (A) Comparison between BCG vaccinated and unvaccinated individuals. (B) Comparison between TST negative and positive individuals. (C) Comparison between male and female. The volcano plots characterize the magnitude (log2 fold change) and the significance (p-values) of antibody response between groups. Values above black dashed lines are statistically different between the groups ($p < 0{\cdot}05$). For adjusted p-values, significant data are shown in red, non-significant differences are in grey.

**Supplemental Figure 5. Antibody functionality for non-TB antigens.** Antibody-dependent neutrophil phagocytosis (ADNP), Antibody-dependent NK cell activation with %CD107a, and Antibody-dependent complement deposition (ADCD) were analyzed in the plasma of children aged 12-60 months, against nonTB antigens: Flu (A) and Tetanus (B). The dot plots show differences in antibody functions between TB and nonTB, as well as ATB and LTBI. Horizontal dashed bars show PBS values. Significance was calculated by using two-sided Mann-Whitney U-test, *$p < 0{\cdot}05$.

**Supplemental Figure 1. Mtb-specific FcR binding in ATB, LTBI and non-TB groups.**

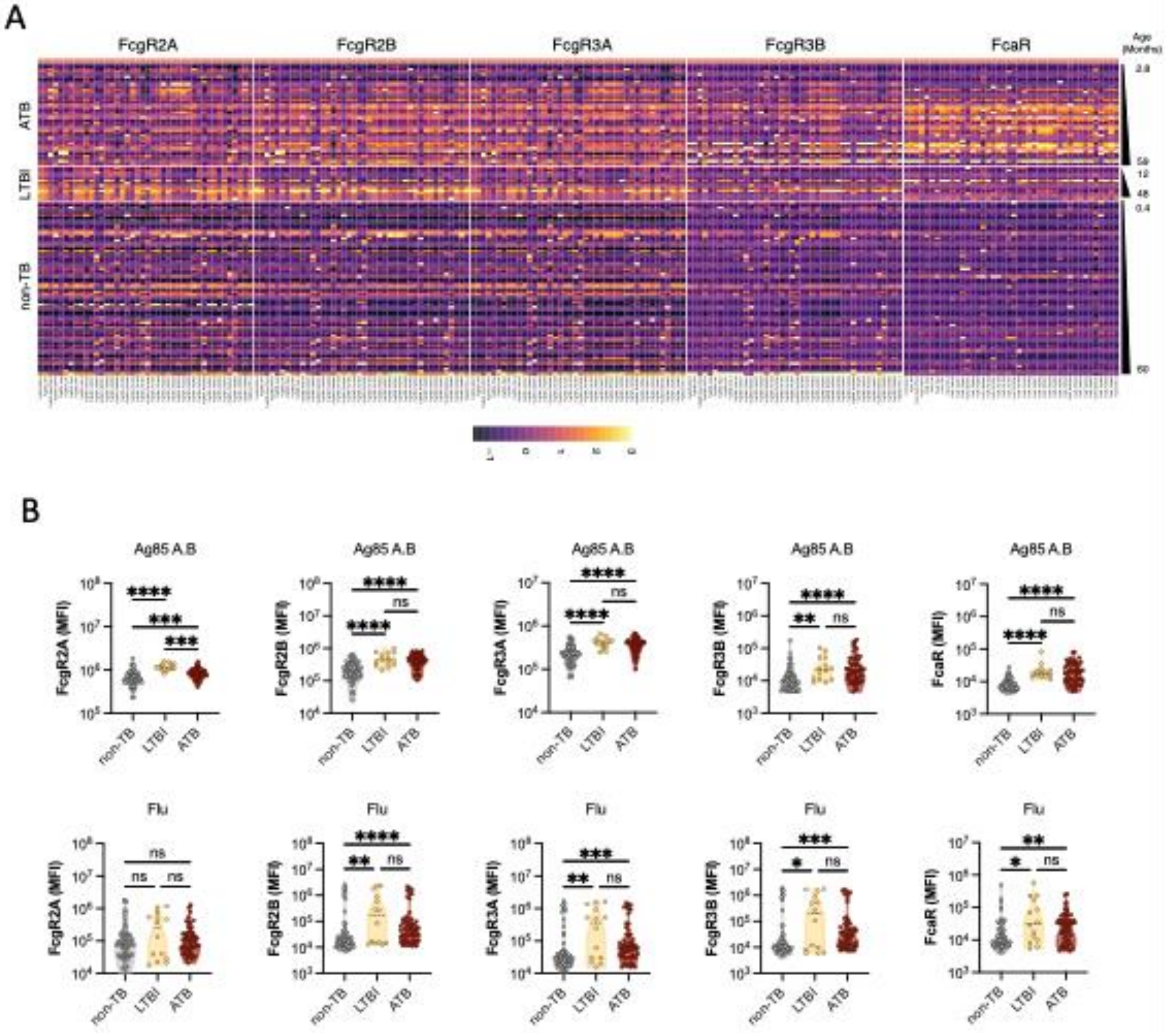

**Supplemental Figure 2. Strong Mtb-specific antibody levels and FcR binding in ATB.**

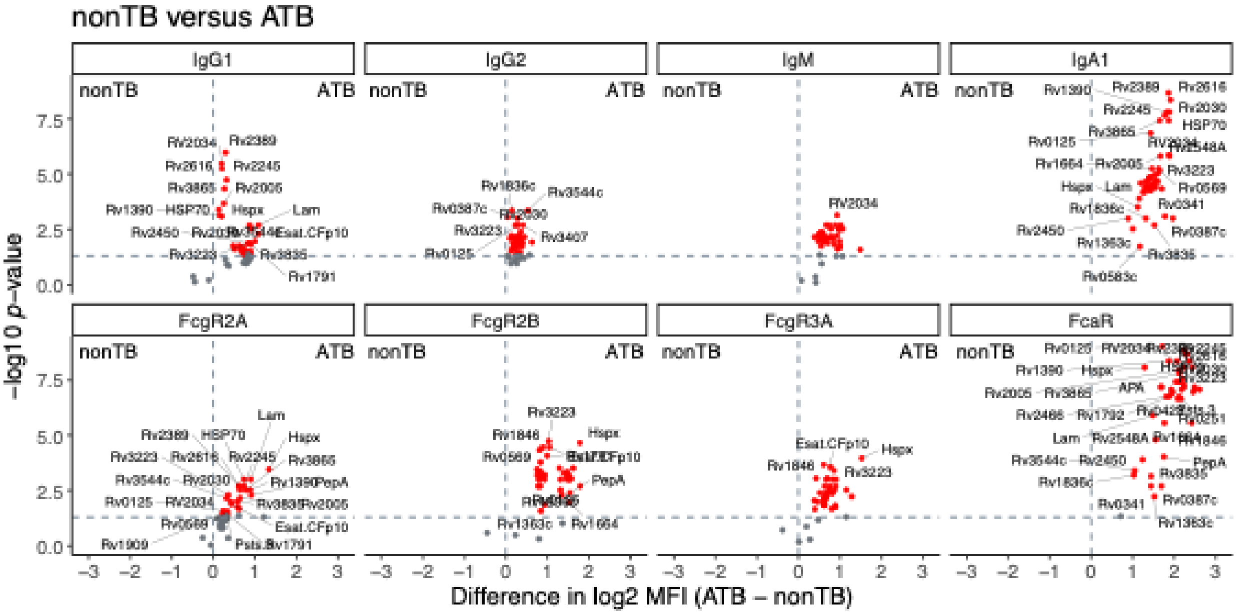

**Supplemental Figure 3. IgG3 and FcgR3B responses against Mtb antigens.**

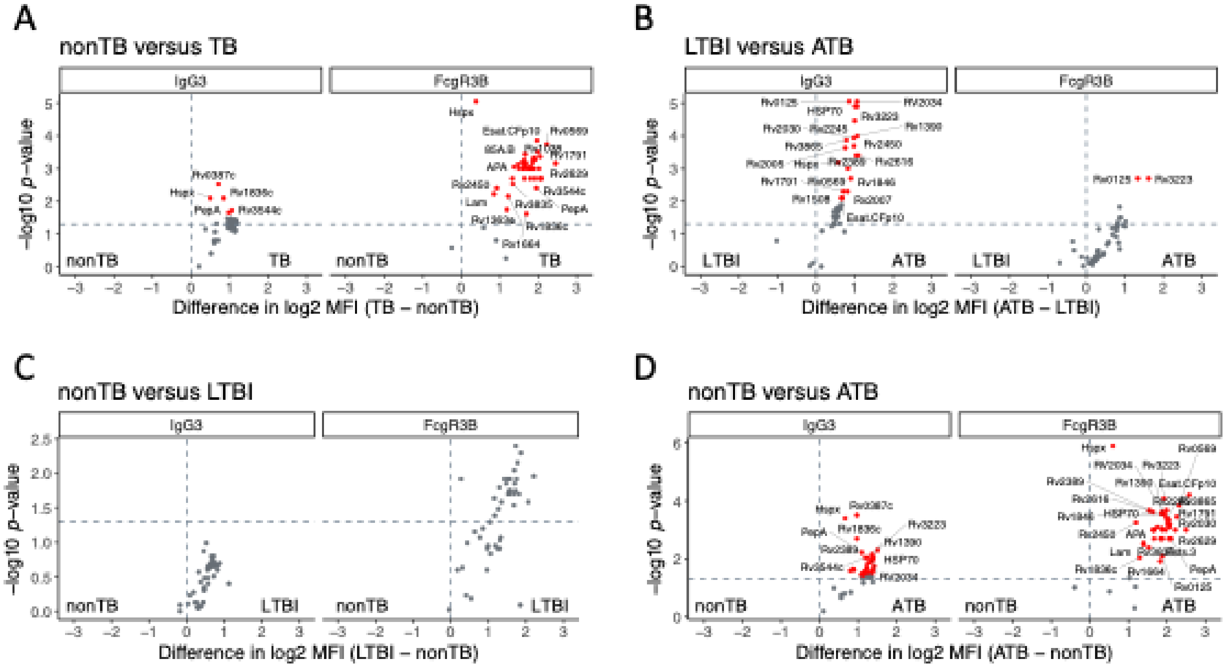

**Supplemental Figure 4. Impact of HIV-infection, BCG vaccination, TST, and sex on antibody response in ATB.**

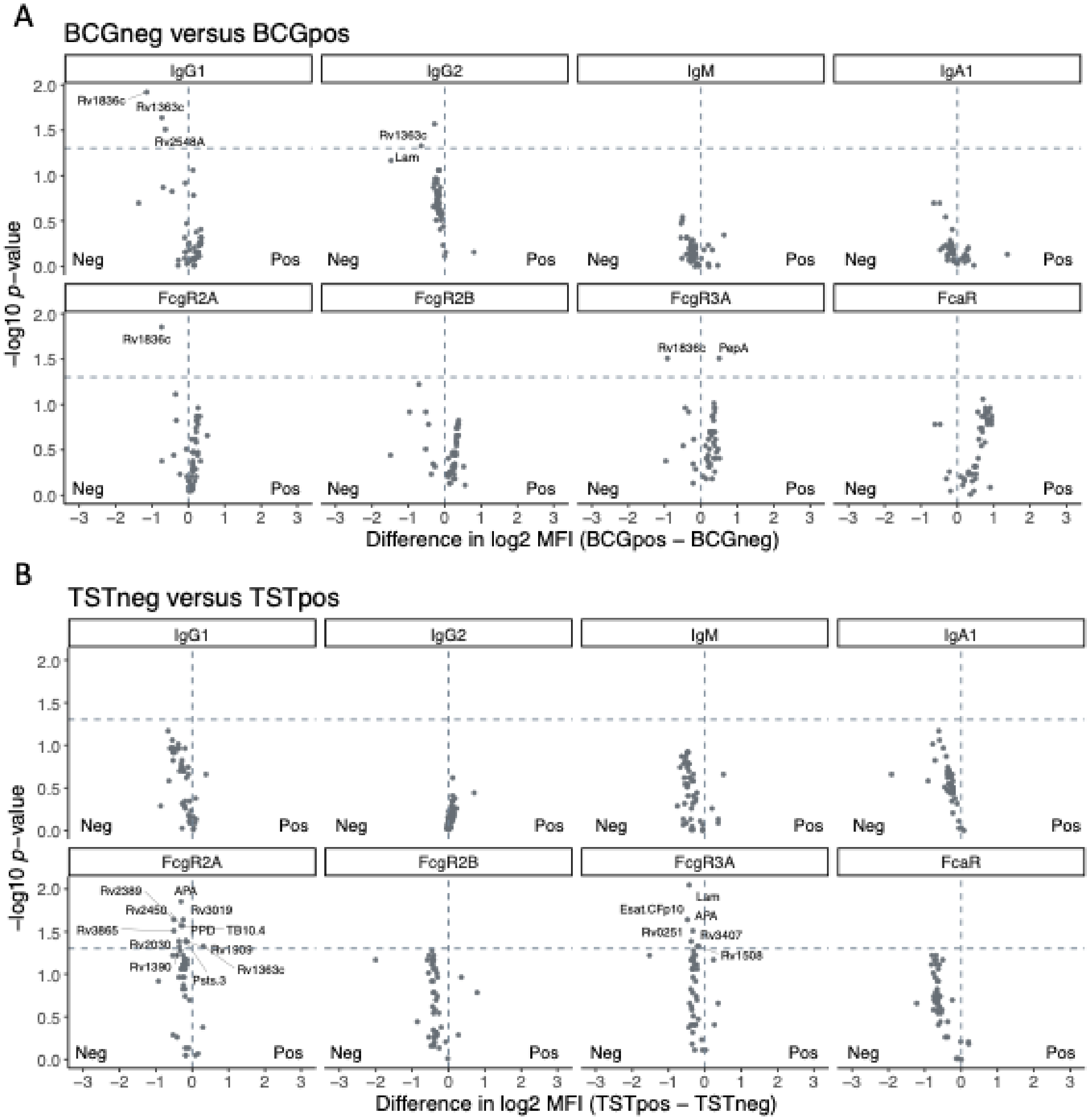

C

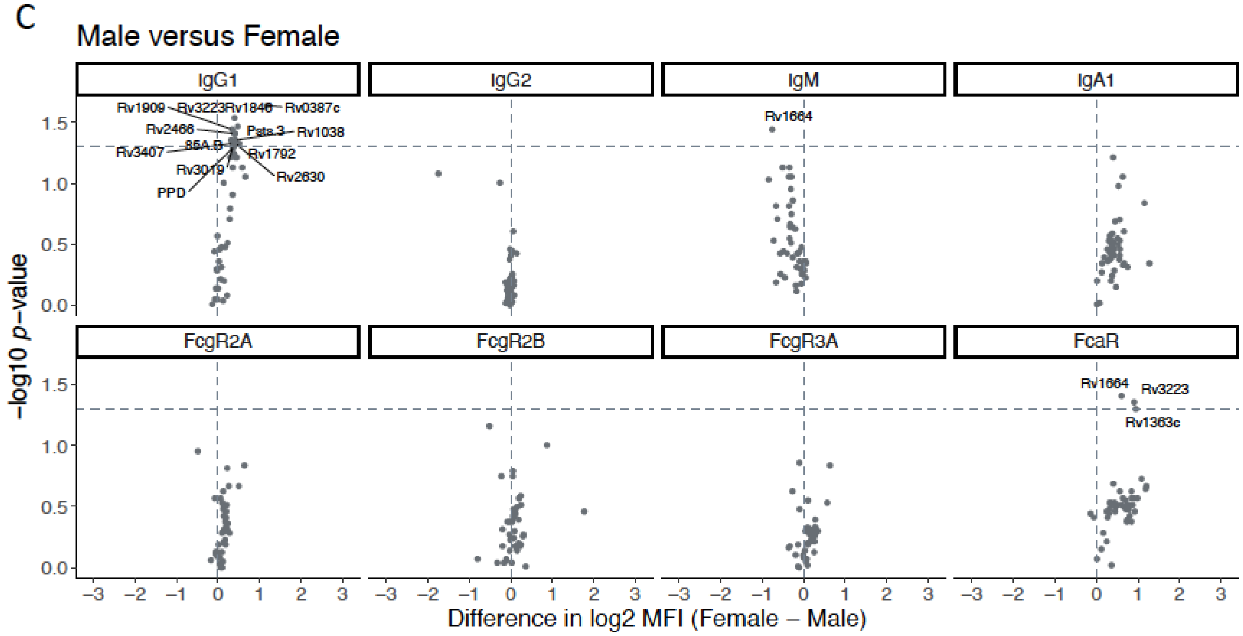

Male versus Female
IgG1
IgG2
IgM
IgA1
FcgR2A
FcgR2B
FcgR3A
FcaR
Rv1909
Rv3223
Rv1846
Rv0387c
Rv2466
Psts.3
Rv1038
85A.B
Rv3407
Rv1792
Rv3019
Rv2630
PPD
Rv1664
Rv1664
Rv3223
Rv1363c
−log10 p−value
Difference in log2 MFI (Female − Male)

**Supplemental Figure 5. Antibody functionality for non-TB antigens**

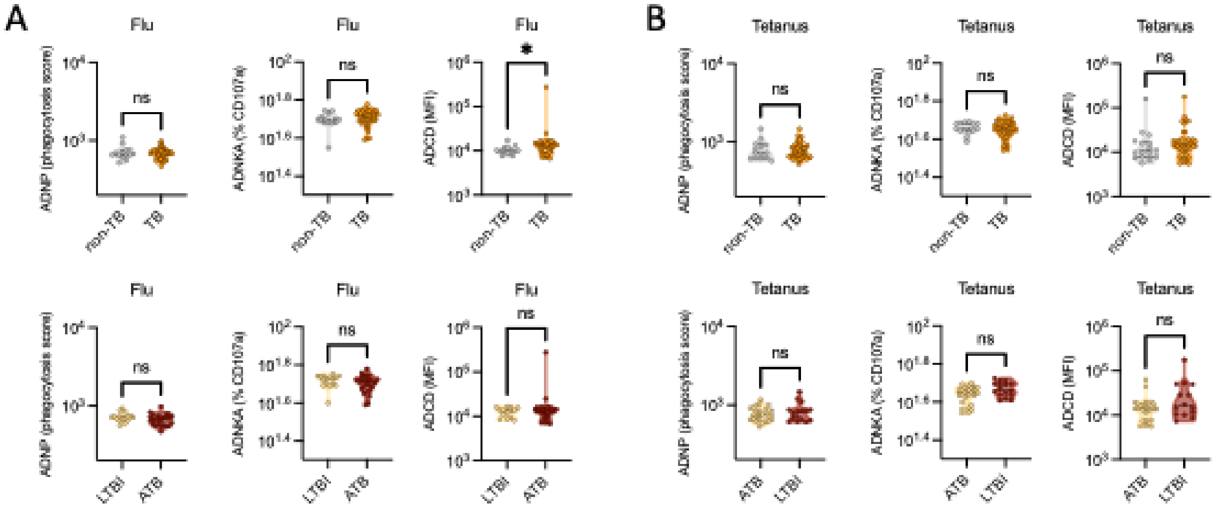
